# Differential Adaptation to a Harsh Granite Outcrop Habitat between Sympatric *Mimulus* Species

**DOI:** 10.1101/091538

**Authors:** Kathleen G. Ferris, John H. Willis

## Abstract

A primary goal in evolutionary biology is to understand which environmental variables and traits drive adaptation to harsh environments. This is difficult since many traits evolve simultaneously as populations or species diverge. Here we investigate the ecological variables and traits that underlie *Mimulus laciniatus*’ adaptation to granite outcrops compared to its sympatric, mesic-adapted progenitor *M. guttatus*. We use fine scale measurements of soil moisture and herbivory to examine differences in selective forces between the species’ habitats, and measure selection on flowering time, flower size, plant height, and leaf shape in a reciprocal transplant using *M. laciniatus x M. guttatus* F_4_ hybrids. We find that differences in drought & herbivory drive survival differences between habitats, that *M. laciniatus* and *M. guttatus* are each better adapted to their native habitat, and differential habitat selection on flowering time, plant stature, and leaf shape. We conclude that while early flowering time, small stature, and lobed leaf shape underlie plant fitness in *M. laciniatus*’ seasonally dry environment, increased plant size is advantageous in a competitive mesic environment replete with herbivores like *M. guttatus*’.

## INTRODUCTION

Closely related species often occupy ecologically disparate habitats. How do different taxa adapt to these new and often stressful environments? What traits evolve in response to novel selection pressures as a species colonizes a new habitat? And which environmental variables act as selective forces? These are exciting questions, especially in sympatric species since adaptation to different habitats can lead to ecological speciation (Mayr 1947; Mayr 1949; Schluter 2001; Coyne & Orr 2004; Rundle & Nosil 2005) or reinforce reproductive isolation between species that have come into secondary contact through selection against migrants or ecologically unfit hybrids (Coyne & Orr 2004). Reciprocal transplant studies have demonstrated differential habitat adaptation between many populations and species, however little is still known about the specific environmental variables and traits behind this adaptation (reviewed in Kawecki & Ebert 2004; Hereford 2009; Salvolainen et al. 2013). This is especially true for closely related species that occupy different habitats. Identifying particular selective agents causing adaptive divergence is a challenge because habitats vary in many ecological variables, and tracking variation in particular ecological variables and their effect on fitness in the field is labor intensive (Wade & Kalisz 1990, MacColl 2011). Understanding the adaptive significance of individual phenotypes is difficult since as species diverge many traits evolve simultaneously due to both adaptive and neutral demographic processes. While many studies have examined natural selection on phenotypes in the wild using the multivariate approach developed by Lande and Arnold (1983; reviewed in Kingsolver et al. 2001), the majority of them measured selection in natural populations where many traits are genetically correlated with each other due to shared demographic history. This extensive genetic correlation among traits makes inferring direct selection on any one phenotype problematic even with a multivariate statistical approach (Hall & Willis 2006). This problem becomes even larger when selection is measured on phenotypic differences between species rather than between populations, since species have a longer history of divergence.

Combining fine scale environmental measurements, controlled crosses between species, and reciprocal transplant experiments can overcome these obstacles. Collecting fine scale ecological data in a reciprocal transplant experiment makes it possible to test whether individual ecological variables are associated with fitness and therefore whether they act as selective agents in a particular habitat (Huber et al. 2004; Garant et al. 2007; Lind & Johansson 2007; Quinn et al. 2009; Weese et al. 2010). In a segregating hybrid population trait associations present in the parental lines are broken up through recombination. This disentangles phenotypes from their species-specific genetic background. Using a segregating hybrid population in a reciprocal transplant experiment therefore allows the measurement of selection on individual traits in the wild (Hall and Willis 2006; Anderson et al. 2011b, Agren et al. 2016).

Plant species are excellent systems for studying differential habitat adaptation. Because of their sessile lifestyle, plants often experience strong divergent selection across heterogeneous environments on a small geographic scale (Kalisz 1986; Schmitt & Antonovics 1986; Stewart & Schoen 1987; Robichaux 1990). Variation in soil moisture is a major driver of plant species’ distribution and diversification (Stebbins 1957; Axelrod 1972). Certain traits have evolved repeatedly in plant species that occur in dry, marginal habitats. Early flowering allows plants to reproduce before the onset of summer drought and has been shown to be adaptive across plant species (Kiang & Hamrick 1987; Fox 1990, Macnair & Gardner 1998; Stanton et al. 2000; Hall & Willis 2006; Willis et al. 2008; Anderson et al. 2012). Plant taxa occupying harsh, marginal habitats also often have self-fertilizing mating systems (Stebbins 1957; Stebbins 1970; Kiang & Hamrick 1987; MacNair & Gardner 1998; Mazer et al. 2010; Wu et al. 2010). Small flower size and rapid floral development are correlated with self-fertilization and may reduce floral tissue transpiration in dry environments (Galen 1999). Additionally, self-fertilization provides reproductive assurance in marginal environments often sparsely populated by pollinators (Stebbins 1970; Piper et al. 1986; Cunningham 2000; Fausto et al. 2001). Diminutive plant stature is adaptive in dry environments because small plants require less water for proper development and therefore are less stressed under drought conditions (Blum and Sullivan 1997; Blum et al. 1997; Pantuwan et al. 2002). Small plants also tend to have more rapid development and flowering time (Thornsberry et al. 2001; Bolmgren and Cowan 2008; Wei et al. 2010). Leaf shape is also hypothesized to be adaptive in dry environments because it affects the gas and heat exchange traits of the plant. Lobed leaves have thinner boundary layers and are cooled more efficiently by convection than round leaves. More efficient convective cooling should decrease the amount of transpiration required when leaves are heated above ambient temperature by direct sunlight (Vogel 1968; Givnish 1978; Schuepp 1993; Nobel 2005). Lobed leaves also have lower hydraulic resistance (*R_leaf_*) and therefore less drought stress prone tissue than round leaves (Nicotra et al. 2011). A thinner boundary layer & lower *R_leaf_* should be advantageous in dry, exposed habitats.

The *Mimulus guttatus* species complex is an excellent system for studying adaptation to harsh marginal habitats because it is a closely related group of wildflowers that are largely inter-fertile and occupy a variety of edaphic environments across Western North America. *Mimulus guttatus*, the putative progenitor of the species complex, occurs in moist seeps and streambeds, but several smaller ranging sympatric species have colonized rapidly drying habitats such as serpentine soils, copper mine tailings, and granite outcrops (MacNair & Gardner 1998; Wu et al. 2007; Ferris et al. 2015; Wright et al. bioRxiv). Granite outcrops are harsh habitats characterized by shallow rocky soils, high light intensity, extreme temperatures, and low soil water retention (Ferris et al. 2014). The Sierra Nevada natives *M. laciniatus* (Figure 1b) and the recently described *M. filicifolius* (Sexton et al. 2013, Ferris et al. 2014) specialize on granitic habitat (Figure 1a). These species grow in thin strips of moss and gravel where few other plant taxa can thrive. Both granite outcrop specialists are sympatric with *Mimulus guttatus* (Figure 1d), which occurs in deeper, densely populated soils in moist seeps and meadows (Figure 1e, Wu et al. 2007). *Mimulus guttatus* can often be found growing in mesic habitat within meters of *M. laciniatus’* granite outcrops (Ferris et al. 2016). *Mimulus laciniatus* and *M. filicifolius* share several traits that are likely adaptive in rocky outcrops that dry rapidly once seasonal snowmelt is gone: early and rapid flowering (Friedman & Willis 2013; Ferris et al. 2016), a self-fertilizing mating system (Fenster & Ritland 1994; Ferris et al. 2014), small plant size, and lobed leaf shape (Sexton et al. 2013, Ferris et al. 2015, Ferris et al. 2016). Lobed leaf shape is unique to rocky outcrop *Mimulus* taxa (Ferris et al.2015), and is seemingly an example of parallel evolution (Ferris et al. 2014), therefore it is also likely adaptive in this marginal habitat. In contrast *M. guttatus* is round-leaved,larger, later-flowering, and predominantly outcrossing (Wu et al. 2007, Ferris et al. 2016).

**Figure 1.**
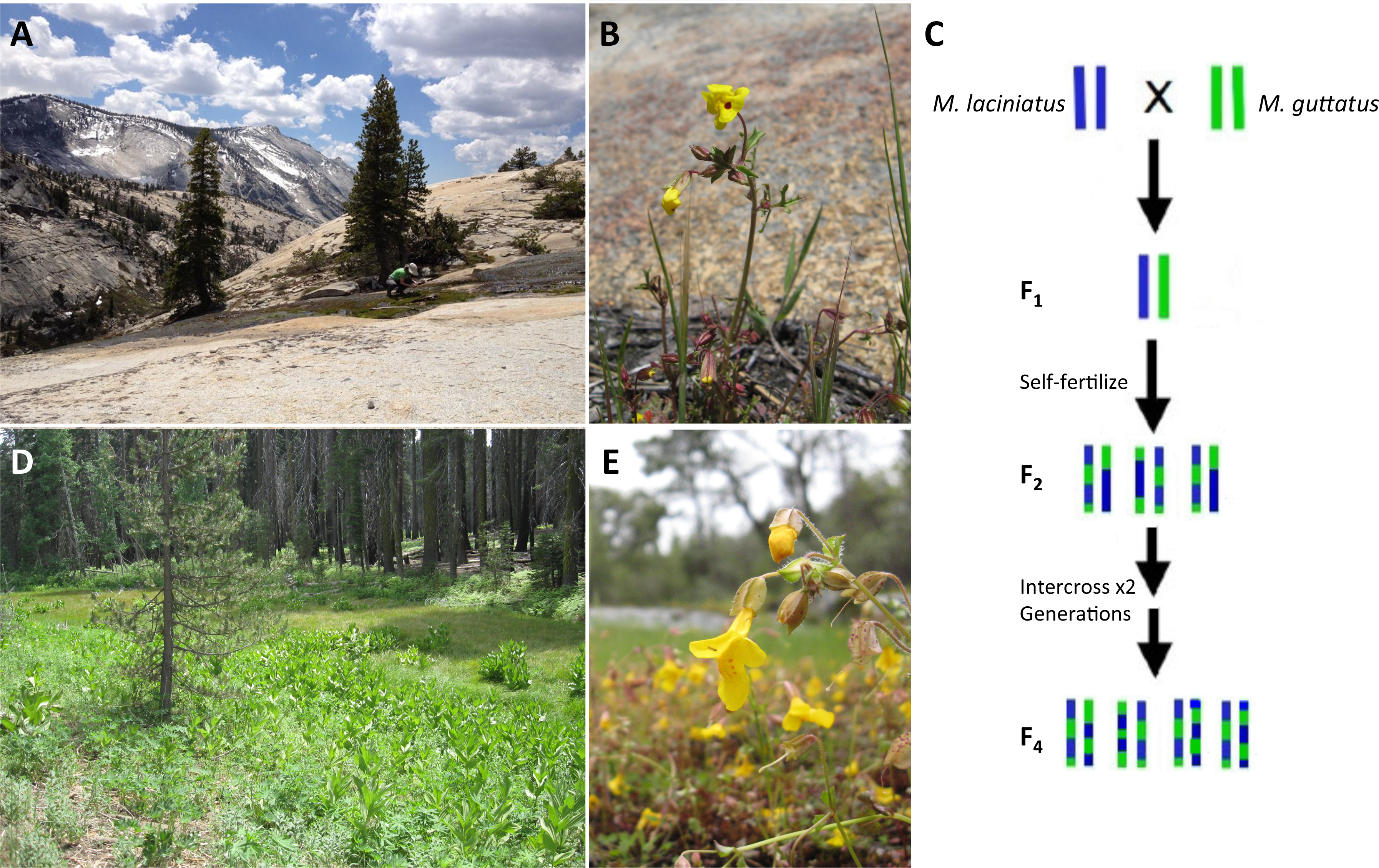
Images of (a) granite outcrop habitat (Granite 1), (b) *M. laciniatus* growing in its native granite habitat, (c) crossing design used to generate the F_4_ hybrid population, (d) meadow habitat (Meadow 2), and (e) *M. guttatus* growing in its native meadow habitat.

**Figure 2.**
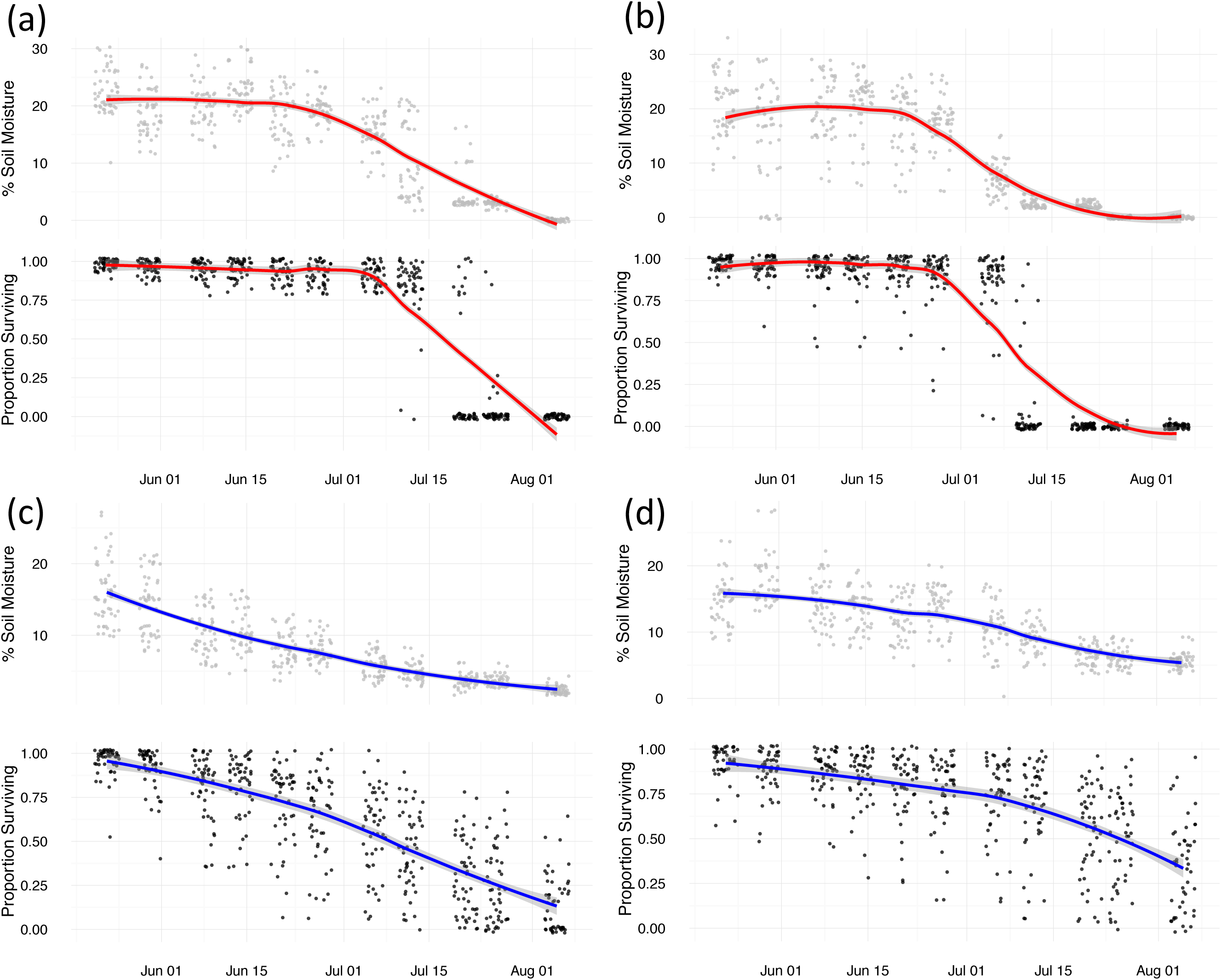
Percent soil moisture (gray) and proportion plants surviving (black) measured weekly over the two month duration of the experiment on a block by block basis across the (a) *M. laciniatus* habitat (red) Granite 1 and (b) Granite 2 sites (c) *M. guttatus* habitat (blue) Meadow 1 and (d) Meadow 2 sites.

In this study we investigate how *M. laciniatus* has adapted to its marginal granite outcrop environment using a reciprocal transplant experiment with *M. laciniatus, M. guttatus*, and a large segregating outbred F_4_ hybrid population planted in replicated native granite and meadow habitats. Trait associations present in the parental species are broken up by recombination and independent assortment in F_4_’s allowing the measurement of selection on individual phenotypes. To understand adaption to *M. laciniatus’* harsh granite outcrop habitats in we asked three questions: (1) What ecological variables act as selective agents in each species’ habitat? (2) Are *M. laciniatus* & *M. guttatus* differentially adapted and if so are there fitness trade-offs? (3) What traits are involved in this adaptation? In a previous survey of microclimatic variation we found that *M. laciniatus’* granite outcrop habitat is significantly drier than adjacent meadow and streambed habitat occupied by *M. guttatus* (Ferris et al. 2014). We also observed lower herbivore density in *M. laciniatus’* granite outcrop habitat than in *M. guttatus’* meadows (K. Ferris personal observation). Soil moisture and herbivory are important factors in plant distribution and diversification (Stebbins 1952; Ehrlich and Raven 1964; Axelrod 1972; Janz 2011). To characterize differences in selective forces between *M. laciniatus* and *M. guttatus’* native habitats we measured variation in these two variables on a fine spatial scale throughout our reciprocal transplant, and tested whether variation in soil moisture or herbivory was associated with survival in each habitat. To test for differential adaption and trade-offs between species we recorded fitness differences between *M. laciniatus* & *M. guttatus* in granite and meadow habitats. To examine individual traits that contribute to *M. laciniatus*’adaptation to a marginal granite outcrop habitat we measured selection in the field on phenotypes that the granite specialists, *M. laciniatus* and *M. filicifolius*, share and which are also likely adaptive in seasonally dry habitats: flowering time, flower size, leaf size, plant size, and a lobed leaf shape in the *M. laciniatus x M. guttatus* F_4_ hybrids. If *M. laciniatus* and *M. guttatus* are differentially adapted to their respective microhabitats and flowering time, flower size, leaf size, plant size, and leaf shape contribute to this adaptation, then we expect there to be divergent selection on these traits between habitats in the direction of species differences.

## MATERIALS AND METHODS

### Construction of the hybrid population

In the summer of 2008 we collected individuals from each parental population, White Wolf (WLF, *M. laciniatus*) and Yosemite Overlook (YVO, *M. guttatus*), along Tioga Road in Yosemite National Park, CA. The parental inbred lines, WLF47 and YVO6, were developed through hand pollinated self-fertilization and single seed descent for 7 and 5 generations respectively. The experimental population of outbred F_4_ hybrids was created by first crossing the parental lines WLF47 and YVO6 to generate F_1_ hybrids, then self-fertilizing F_1_’s to generate a large F_2_ population. Two hundred F_2_’s were randomly paired and reciprocally crossed to generate 200 maternal families. Then 200 F_3_’s were randomly paired and reciprocally crossed (Figure 1c). We pooled ˜30 seeds from each of the 200 maternal F_3_’s to create one large outbred F_4_ family. F_4_ seeds were then randomly distributed throughout experimental blocks.

### Reciprocal transplant experiment

In April of 2013, 400 *M. laciniatus* (WLF47), 400 *M. guttatus* (YVO6), and 3000 F_4_’s were planted in open flats of soil in the UC Davis greenhouses. Seeds were cold stratified at 4°C for 10 days, then left in the greenhouse to germinate for one week. At the cotyledon stage seedlings were transplanted one inch apart into 50 randomized blocks of 19 plants each (2 WLF47, 2 YVO6, 15 F_4_) at each of four field sites along Tioga Road in Yosemite NP, CA. The two *M. laciniatus* habitat sites, Olmstead Point (Granite 1, Figure 1a) and Yosemite Creek (Granite 2), are undisturbed granite outcrops with native *M. laciniatus* growing on moss at elevations of 8,500 and 7,500 feet respectively. The *M. guttatus* sites, Little Meadow (Meadow 1) and Crane Flat (Meadow 2, Figure 1e), are undisturbed meadows with native *M. guttatus* growing near a standing seep. They occur at 6,200 and 6,000 feet respectively. Sites were chosen to maximize the similarity between the developmental stages of transplanted and native plants of each species.

Seedlings that died within a week of transplant were replaced so that transplant shock would not affect survivorship. Experimental plants were censused every other day May - August 2013 for survival and flowering time. On the day of first flower, flowering time, corolla width, and plant height were recorded. Morphological measurements were taken with a small metal ruler in millimeters. Leaves were not collected for leaf shape measurement until plants began to senesce to avoid damaging plants and imposing selection before they set seed. Leaf area and degree of lobing were determined by digitally scanning leaves and performing convex hull analysis in ImageJ as described previously (Ferris et al. 2015). Briefly, the convex hull analysis consists of comparing the area of each leaf’s convex hull (the shape created by connecting the outermost points of a leaf) to the leaf’s true area and dividing this difference in area by the convex hull area to control for size. Fruit number was counted after senescence as a measure of lifetime fitness.

### Measuring environmental agents of selection

We collected fine scale soil moisture and herbivore damage data in each block across each transplant site. This allowed us to test whether these environmental variables are associated with plant fitness differences on a fine spatial and temporal scale within and between habitat types. We used survival as our measure of fitness in this analysis because we had data on survival in every block in each site giving us greater spatial variation in survival than in fruit number since only a subset of experimental blocks produced fruit. Percent soil moisture was measured weekly in each experimental block May through August using a Decagon soil moisture probe. We measured herbivory in each block at the time of plant senescence by recording the presence or absence of damage on the first true leaf.

## ANALYSES

### Selective environment

To determine whether drought and herbivory contributed to the difference in species’ selective environments we measured percent soil moisture and leaf damage in each experimental block of our reciprocal transplant experiment. We tested if soil moisture and leaf damage affected plant fitness across habitats with a linear mixed effects model where soil moisture was treated as a repeated measure using the nlme package in R (R Core Development Team 2008). In our mixed effects model survival was the dependent variable with habitat type, soil moisture, leaf damage, and time as fixed independent variables, and block as a random effect nested within site, nested within habitat. To determine whether different environmental variables were associated with plant survival in granite vs. meadow sites we tested for interactions between our environmental variables and habitat type. We also ran separate mixed effects models on the survival data from each habitat. We determined the best-fit model using AIC model selection criteria (Zuur et al. 2009) with the model selection R-package MuMin (Bartoń 2013). To determine whether there was an absolute difference in herbivore damage between habitat types regardless of its effect on survival, we performed a t-test. If soil moisture and herbivory are selective agents in either granite or meadow habitats then we expect a strong association between the environmental variable and survival within a transplant site. If either variable contributes to differential selection between habitat types, then we expect a stronger association in one habitat over the other.

### Habitat adaptation and trait differences

To test for differential habitat adaptation between *M. laciniatus* and *M. guttatus* we performed two-tailed t-tests to determine whether mean survival to flowering and fruit number significantly differed between *M. laciniatus* and *M. guttatus* by habitat. We also conducted a logistic regression with survival to flowering as the dependent variable and species, habitat, and their interaction as independent variables. If *M. laciniatus* and *M. guttatus* are differentially adapted then we expect each species to have a fitness advantage in its native habitat. If there is a species trade-off between habitats we expect to see an interaction between habitat type and species in our regression. Based on our phenotypic results we also decided to look for species differences in the degree and direction of phenotypic plasticity in each trait. To detect phenotypic plasticity in flowering time, flower size, plant height, leaf area, and leaf shape we conducted factorial ANOVAs with habitat (Granite vs. Meadow) & genotypic class (*M. laciniatus, M. guttatus*, & F_4_) as independent variables. To detect genotype by environment (GxE) interactions we tested for interactions between habitat and genotypic class. To correct for multiple testing we used a Bonferroni correction (α = 0.01). If there is a difference in the degree or direction of phenotypic plasticity between species, then we should see a significant interaction between genotypic class and habitat (GxE). Phenotypic correlations were measured among traits for each habitat type using a restricted maximum likelihood model (REML) in JMP (JMP®, Version *10*. SAS Institute Inc., Cary, NC, 1989-2007).

### Phenotypic selection analysis

To measure the strength of selection on individual phenotypes we conducted linear and quadratic selection analysis (Lande & Arnold 1983; Mitchell-Olds & Shaw 1987). We used lifetime fitness (fruit number) of plants that survived to flower as our fitness measure in our selection analysis because we only have phenotypic values of plants that survived to flower. Fecundity, fruit number in our case, is also the best measure of fitness for an annual plant. We used a generalized linear mixed model to regress lifetime fitness simultaneously on flowering time, corolla width, leaf area, plant height, and leaf shape in our F_4_ population. To detect differential habitat selection we tested for interactions between trait and habitat type. We also measured the strength of selection on each phenotype in each habitat by performing a regression on data from each habit type separately. Block and site were included in the model as random effects where block was nested within site, which was nested within habitat. The strength of selection on each trait in each habitat was measured with linear (β’) and quadratic (γ) regression coefficients. All phenotypes were standardized to a mean of 0 and standard deviation of 1 to enable comparison of traits measured in different units and to values from other phenotypic selection studies (Lande & Arnold 1983).

To control for excess zeros in our lifetime fitness measure we broke our data up into two distinct analyses: (1) a binomial logistic regression on all plants the flowered whether or not they produced fruits, and (2) a truncated Poisson regression on only plants that did produce fruits (as in Anderson et al. 2012, Anderson et al. 2015) using the R package glmmADMB (Skaug et al. 2006). The binomial distributed logistic regression determines whether a trait was associated with the probability of a plant setting seed or not. The truncated Poisson regression on the count data for plants that did fruit tested whether phenotypes were associated with the number of fruits a plant produced (Ridout et al. 1998). Model selection was performed using AIC selection criteria (Zuur et al.2009) with the model selection R-package MuMin (Bartoń 2013). Phenotypes and interactions included in the top model were considered to be under selection. If the phenotypes we have measured contribute to differential habitat adaptation between these species we expect divergent selection in the direction of the native species trait value. For example, if flowering time is involved in differential adaptation we would expect there to be selection for earlier flowering in *M. laciniatus*’ granite outcrops, but for later flowering in *M. guttatus*’ meadow habitat.

## RESULTS

### Drought and herbivory are differentially associated with survival differences between habitats

When the environmental data from all sites was analyzed together our best-fit mixed effects model included both soil moisture and leaf damage, indicating that both significantly affected plant survival across habitats (Table 1). Our best model also included an interaction between soil moisture & habitat and a three-way interaction between soil moisture, habitat, and time indicating that soil moisture affects survival differently in each habitat over time (Table 1). Specifically, while soil moisture is in fact higher in granite outcrops at the beginning of the growing season, it then decays rapidly to effectively zero over the course of approximately one week mid-summer (Figure 2a). In contrast, soil moisture in the meadow habitat decays in a gradual linear fashion over the course of the entire season (Figure 2b). Survival tracks soil moisture with a slight delay in both granite and meadow habitats (Figure 2c&d). When data from each habitat type was analyzed separately soil moisture was associated with survival in both, but leaf damage only affected survival in meadows (Table 1). There was also significantly more herbivore damage, and therefore more herbivore pressure, in the meadows than in granite (p-value < 0.001, Figure 3).

**Figure 3.**
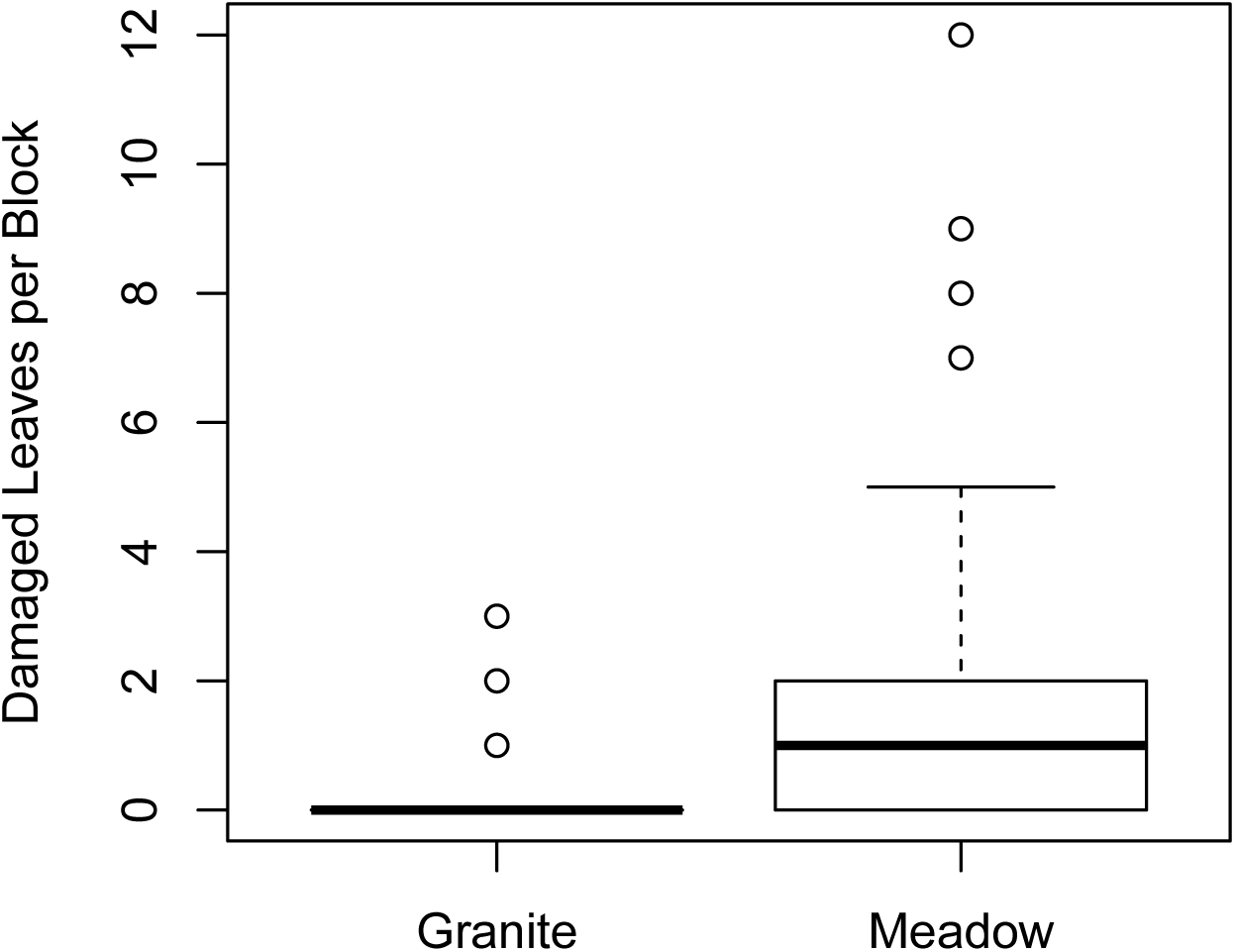
A boxplot of the number of herbivore damaged leaves per experimental block at the time of plant senescence in granite vs. meadow habitat.

**Table 1.** Results of the linear mixed effects model testing how environmental variables affected survival across sites and time given as the equations with the highest AIC criteria. In the model description SM = Soil Moisture, LD = Leaf Damage, T=Time, and H=Habitat. “k” is the number of parameters in the model, “Log(L)” is the log likelihood, “AIC” score, “Δi” is the difference between the AIC score of the current & top models, “wi” is the model weight, Marginal R^2^ is the contribution of the fixed effects, Conditional R^2^ is the total contribution of both random and fixed effects.

| Model | k | Log(L) | AIC | Δi | wi | Marginal R<sup>2</sup> | Conditional R<sup>2</sup> |
| --- | --- | --- | --- | --- | --- | --- | --- |
| Survival Total ~ SM + LD + T + H + SM*T + SM*H + LD*SM + SM*T*H | 10 | 977.1 | -1912 | 0 | 1 | 0.802 | 0.833 |
| Survival Granite ~ SM + T + SM*T | 5 | 399.6 | -774.8 | 0 | 1 | 0.853 | 0.856 |
| Survival Meadow ~ SM + T + LD + SM*T | 6 | 962.4 | -1896 | 0 | 0.607 | 0.627 | 0.692 |

### M. laciniatus & M. guttatus are differentially adapted and phenotypically plastic

There was strong viability selection in each site with the total proportion surviving to flower ranging from 0.18-0.57 (G1 = 0.45, G2 = 0.22, M1 = 0.18, M2 = 0.57). *M. laciniatus* had significantly higher survival than *M. guttatus* in its native granite (0.23 > 0.12, p-value < 0.0001), but the species had similar survival in *M. guttatus’* meadows (*M. guttatus* **=** 0.09, *M. laciniatus* = 0.10, p-value = 0.72, Figure 4a). However, a fitness trade-off in fecundity was indicated by crossing reaction norms (Figure 4b). *M. laciniatus* has significantly higher mean fruit number in granite habitat (mean = 0.48, SE = 0.05) than *M. guttatus* (mean = 0.10, SE = 0.022, p-value <0.0001), while *M. guttatus* has marginally significantly higher mean fruit number than *M. laciniatus* in its meadows (*M. guttatus* = 0.05, SE = 0.022, *M. laciniatus* = 0.007, SE = 0.007, p-value = 0.055).In addition, our logistic regression of survival to flowering detected a significant interaction between habitat and species indicating that parental lines are differentially adapted (Table S1).

**Figure 4.**
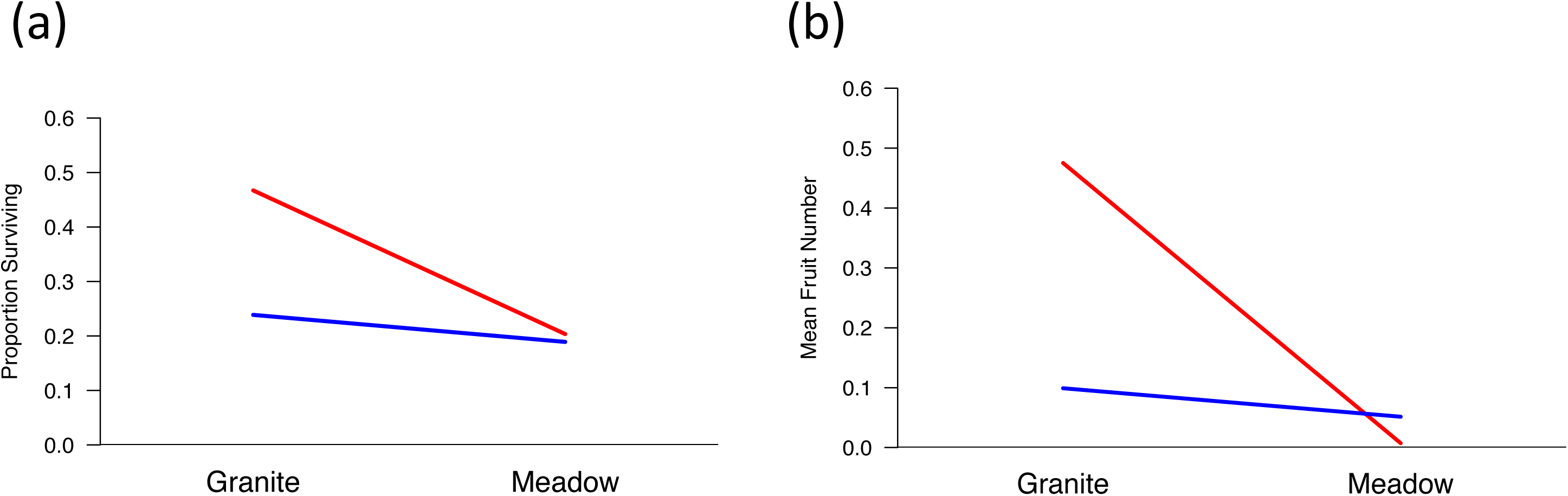
Reaction norm plots of (a) mean survival & (b) fecundity of *M. laciniatus* (red) and *M. guttatus* (blue) parental inbred lines in granite (*M. laciniatus*) and meadow (*M. guttatus*) habitats.

To understand how ecologically important traits varied between habitats and species we calculated the mean trait value and standard error for F_4_’s, *M. laciniatus*, and *M. guttatus* in each habitat and transplant site. In granite habitat phenotypic differences between genotypes are in the expected direction with *M. laciniatus* flowering earlier (by 3.29 days), being smaller flowered (by 2.88 mm) and leaved (by 51.84 mp), shorter (by 2.64), and more lobed (by 0.07) than *M. guttatus* (Table 2). In the meadow habitat the direction of trait differences was the same, except that *M. laciniatus* flowered 3.5 days later than *M. guttatus* (Table 2).

We saw plasticity in our focal traits between habitats and decided to test whether there was significant phenotypic plasticity and genetic variation for plasticity between *M. laciniatus* and *M. guttatus*. There was significant phenotypic plasticity in all traits except leaf area, significant GxE in flowering time (p-value = 0.007), and marginally significant GxE in plant height (p-value = 0.022). Within each genotypic class (F_4_, *M. laciniatus, M. guttatus*) plants flowered earlier (8.57, 8.8, 2.05 days, p-value < 0.001) and were shorter (4.63, 17.94, 43.23 mm, p-value < 0.001) in *M. laciniatus’* granite habitats than in *M. guttatus’* meadows (Table 2, Figure 5). Interestingly *M. laciniatus* was significantly more plastic in flowering time, but less plastic in height than *M. guttatus* (p-value = 0.007, 0.022, Figure 5&c). F_4_’s and *M. laciniatus* had larger flowers (2.2 mm) in granite, while *M. guttatus* had larger flowers (1 mm) in meadows (p-value < 0.001). *M. laciniatus* & F_4_’s showed a slight, but significant (p-value = 0.002) increase in leaf lobing in the granite habitat (Tables 1, Figure 5e).

**Table 2.** Phenotypic trait means and standard errors calculated for day of first flower (days), plant height (mm), flower width (mm), leaf are (megapixels), and leaf shape *M*. *laciniatus’s* granite outcrop versus *M. guttatus’s* meadow habitat type in F_4_’s, *M*. *laciniatus*, and *M. guttatus*. Significant phenotypic plasticity in a trait is indicated by “*” and significance codes are as follows: p-value < 0.001 = ***, 0.01= **,0.05 =*.

| Plant | Trait | Mean Granite | SE Granite | Mean Meadow | SE Meadow |
| --- | --- | --- | --- | --- | --- |
| *F4* | Days to 1st Flower | 77.85*** | 0.263 | 86.42*** | 0.774 |
|  | Height | 40.55*** | 0.789 | 45.18*** | 1.764 |
|  | Flower Width | 8.45*** | 0.131 | 6.292*** | 0.246 |
|  | Leaf Area | 2009 | 157.6 | 3203 | 281.2 |
|  | Leaf Shape | 0.13** | 0.002 | 0.12** | 0.004 |
| *M. laciniatus* | Days to 1st Flower | 76.96*** | 0.505 | 85.78*** | 1.327 |
|  | Height | 28.59*** | 1.089 | 46.53*** | 4.766 |
|  | Flower Width | 6.075*** | 0.253 | 3.875*** | 0.467 |
|  | Leaf Area | 1068 | 144.0 | 1307 | 248.2 |
|  | Leaf Shape | 0.18** | 0.011 | 0.17** | 0.018 |
| *M. guttatus* | Days to 1st Flower | 80.25*** | 0.743 | 82.3*** | 1.739 |
|  | Height | 31.23*** | 1.667 | 74.46*** | 7.032 |
|  | Flower Width | 8.8*** | 0.42 | 9.835*** | 0.473 |
|  | Leaf Area | 1120 | 103.1 | 3524 | 647.1 |
|  | Leaf Shape | 0.11 | 0.006 | 0.11 | 0.01 |

**Figure 5.**
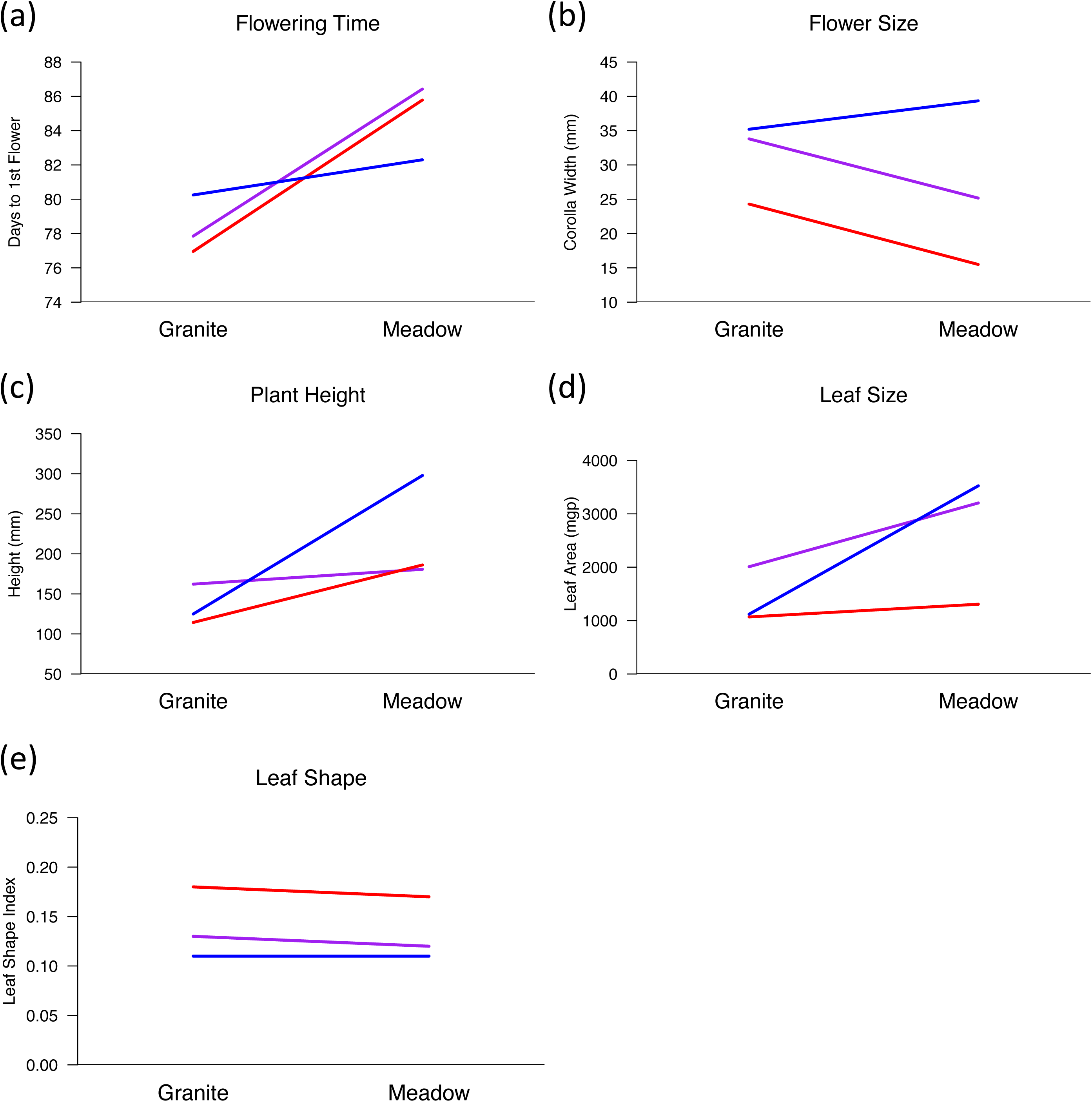
Reaction norm plots demonstrating phenotypic plasticity between granite and meadow habitats for (a) mean flowering time (b) mean corolla width (c) mean plant height (d) mean leaf area, and (e) leaf shape for each genotypic class: *M. laciniatus* (red), *M. guttatus* (blue), and F_4_’s (purple).

We examined phenotypic correlations among traits in the F_4’_s (Table 3). Phenotypic correlations in a segregating hybrid population can provide insight concerning the genetic correlations among traits. In both habitats leaf area, flower size, and height at first flower were highly positively correlated (r > 0.5), while leaf area and shape are slightly positively correlated (granite r = 0.079, meadow r = 0.136). This suggests that there is a genetic correlation among these traits in both habitats. Flowering time was uncorrelated with other traits in granite (Table 3), but positively correlated with height (r = 0.110) and leaf area (r = 0.128) in meadows. This difference suggests that the genetic architecture of flowering time could differ between habitats.

**Table 3.** Phenotypic correlation matrix from REML analysis among all traits at granite vs. meadow sites separately. The granite habitat correlations are in the bottom matrix in black and the meadow habitat correlations are on the top in gray.

|  | Days to Flowering | Stem Length | Flower Width | Leaf Area | Shape |
| --- | --- | --- | --- | --- | --- |
| Days to Flowering | **1** | 0.11 | 0.023 | 0.128 | -0.058 |
| Stem Length | -0.073 | **1** | 0.557 | 0.534 | 0.010 |
| Flower Width | 0.047 | 0.677 | **1** | 0.305 | 0.0002 |
| Leaf Area | -0.088 | 0.525 | 0.34 | **1** | 0.135 |
| Shape | -0.052 | 0.046 | 0.08 | 0.08 | **1** |

### Early flowering time is adaptive in granite outcrops, while being large is adaptive in meadows

To assess selection on individual phenotypic traits in each habitat we performed linear and quadratic selection analysis in our outbred F_4_ population using a combination of binomial logistic and truncated Poisson generalized linear mixed models (GLMM). In the binomial logistic model we tested whether phenotypic values were correlated with the probability that an individual F_4_ produced fruit or not. Our best-fit binomial model included linear selection gradients on flower size, plant height, leaf shape, flowering time, a negative quadratic selection gradient on flowering time, an interaction between flowering time and plant height, and an interaction between habitat type and flowering time (Table 4). In the granite habitat we found directional selection for larger flower size, strong selection for smaller plant size and earlier flowering time, and weak selection for more highly lobed leaves (Table 5). We also found a signature of stabilizing selection on flowering time in granite indicated by a negative quadratic selection gradient (Table 5; Lande & Arnold 1983). In the meadow habitat we saw selection for earlier flowering time, smaller flowers, less lobed leaves, and strong selection for larger plant size (Table 5). We also detected a signature of stabilizing selection on plant height in meadows. The interaction between flowering time and plant height in our top binomial model suggests correlated selection on these two traits and a potential functional correlation (Lande & Arnold 1983). Selection for earlier flowering time in granite was significantly stronger than selection for early flowering in the meadow habitat indicated by the interaction between habitat and flowering time in our top binomial model (Tables 4&5). We list all binomial distributed generalized linear mixed models testing for phenotypic selection in the supplementary materials (Table S2).

**Table 4.** A list of the top phenotypic selection generalized linear mixed models based on AIC score. For each analysis we report the complete model, k which is the number of parameters, Log(L) which is the likelihood score, AIC score, i which is the difference between that model‘s AIC and the top model‘s AIC, and wi which is the model weight. FT stands for flowering time, FW stands for flower width, HT stands for height, LA stands for leaf area, LS stands for leaf shape, and HB stands for habitat.

| Model | k | Log(L) | AIC | $\Delta_i$ | $w_i$ |
| --- | --- | --- | --- | --- | --- |
| <u>Binomial</u> |  |  |  |  |  |
| Total Fitness ~ FT + FW + HT + LA + LS + HB + FT<sup>2</sup> + FT*HB + FT*HT | 11 | -329.388 | 687.517 | 0 | 0.420 |
| Lifetime Fitness Granite ~ FT + FW + HT + LS + FT<sup>2</sup> + FT*HT | 10 | -238.845 | 498.270 | 0 | 0.886 |
| Lifetime Fitness Meadow ~ FT + FW + HT + LS + HT<sup>2</sup> + LS*HT | 10 | -82.893 | 187.002 | 0 | 0.372 |
| <u>Truncated Poisson</u> |  |  |  |  |  |
| Lifetime Fitness ~ FT + FW + HT + LS + HB + FT*HB + LS*HB + LS*HT | 12 | -123.617 | 272.659 | 0 | 0.293 |
| Lifetime Fitness Granite ~ FT + FW + HT + LS + LS*HT | 8 | -101.696 | 220.206 | 0 | 0.206 |
| Lifetime Fitness Meadow ~ FT + FW + HT | 6 | -21.426 | 57.006 | 0 | 0.246 |

**Table 5.** Results of phenotypic selection analysis on flowering time, flower width, plant height, leaf area, and leaf shape. This table contains the regression coefficients from the binomially distributed logistic generalized linear mixed model (glmm) with the probability that the plant set fruit as the response variable. β represents the linear selection gradient and γ represents the quadratic selection gradient for each phenotypic trait in each habitat type. Significant codes are as follows: p-value < 0.001 = ***, 0.01= **,0.05 =*.

| Trait | $\beta$ Granite | $\beta$ Meadow | $\gamma$ Granite | $\gamma$ Meadow |
| --- | --- | --- | --- | --- |
| Flowering Time | -0.957 | -0.216 | -0.801 | -- |
| Flower Width | 0.331 | -0.279 | -- | -- |
| Height | -1.256 | 1.329 | -- | -0.55 |
| Leaf Shape | 0.066 | -0.273 | -- | -- |
| LS*Height | -- | -- | -0.096 | 0.062 |

In the truncated Poisson regression, which tested for associations between phenotypic value and fruit number among plants that did set seed, our best-fit model included linear selection gradients for flowering time, flower size, plant height, leaf shape, an interaction term for leaf shape and plant height, and interactions between flowering time and habitat and leaf shape and habitat (Table 4). There was directional selection for earlier flowering, larger flowers, taller plants, and more highly lobed leaves in the granite habitat (Table 6). In the meadow habitat we found weaker selection for earlier flowering time, selection for larger flowers, and stronger selection for taller plants (Table 6). We found interactions between habitat type and flowering time and habitat and leaf shape in our top Poisson model (Table 6) indicating that there is differential habitat selection on these two phenotypes. In our second to top model we also saw an interaction between habitat and plant height (Table S3) suggesting that height may also be under differential selection. We list all Poisson distributed generalized linear mixed models testing for phenotypic selection in the supplementary materials (Table S3). The interaction between leaf shape and plant height in our top truncated Poisson model suggests correlated selection on these two traits and a potential functional correlation between them (Lande & Arnold 1983). Leaf area was not included in either the top binomial or Poisson models and therefore we do not include any selection gradient values for leaf area. We recognize that selection gradient values could be due to selection on unmeasured correlated characters, although because we used a segregating hybrid population there should be fewer genetically correlated phenotypes.

**Table 6.** Results of phenotypic selection analysis on flowering time, flower width, plant height, leaf area, and leaf shape. This table contains the regression coefficients from the truncated Poisson distributed generalized linear mixed model on fruit count of plants that did produce fruit. β represents the linear selection gradient and γ represents the quadratic selection gradient for each phenotypic trait in each habitat type and population. Significance codes are as follows: p-value < 0.001 = ***, 0.01= **,0.05 =*.

| Trait | $\beta$ Granite | $\beta$ Meadow | $\gamma$ Granite | $\gamma$ Meadow |
| --- | --- | --- | --- | --- |
| Flowering Time | -1.535 | -0.228 | -- | -- |
| Flower Width | 0.47 | 0.484 | -- | -- |
| Height | 0.555 | 0.999 | -- | -- |
| Leaf Shape | 0.135 | -- | -- | -- |
| LS*Height | -- | -- | -0.479 | -- |

## DISCUSSION

Since the pioneering reciprocal transplant experiments of Clausen, Keck, and Hiesey (1941) differential habitat adaptation has been repeatedly demonstrated in plants (Hereford 2009). However the selective forces and traits driving differential adaptation are still poorly understood (Salvolainen et al. 2013). This is especially true of differentially adapted species. When closely related species are sympatric adaptation to different environments can increase reproductive isolation between them (Coyne & Orr 2004) and promote species coexistence (Wright 2002). To test the adaptive significance of early flowering time, small flower size, small plant size, and lobed leaf shape in *M. laciniatus’* marginal rocky outcrop habitat we performed a reciprocal transplant with *M. laciniatus, M. guttatus*, and outbred F_4_’s. We collected fine scale soil moisture and herbivory data, measured fitness of the parental species, and selection on phenotypes in F_4_’s in each habitat. Drought and herbivory are differentially associated with survival in each habitat making them likely agents of divergent selection. *Mimulus laciniatus* is better adapted to granite than its close relative and progenitor *M. guttatus*, and this adaptation is underlain by differential selection on flowering time, plant size, and leaf shape between habitats.

### Soil moisture & herbivory are likely selective agents

Determining the selective forces that drive adaptation to different in environments is an important, but challenging component of the study of adaptive evolution (Wade & Kalisz 1990). It is a challenge because many environmental variables differ between any two habitats and any number of them may affect fitness (MacColl 2011). To test the significance of a particular ecological variable it is necessary to isolate the impact of that variable on fitness either through (1) environmental manipulation (Bright 1998; Stanton et al. 2000; Weijshede et al. 2008; Campitelli & Stinchcombe 2013a), or (2) through fine scale environmental measurements within and between habitat types (Huber et al. 2004; Garant et al. 2007; Lind & Johansson 2007; Quinn et al. 2009; Weese et al. 2010). We have assessed the selective environments of granite and meadow habitats through the latter approach. We measured soil moisture and herbivory, which we had prior reason to believe were selective factors in these habitats (Peterson et al. 2013, Ferris et al. 2014, K. Ferris personal observation), on a fine spatial scale throughout each reciprocal transplant site so that we could determine whether variation in these factors was associated with variation in survival within each habitat type. We also measured soil moisture on a fine temporal scale. Then we examined whether there was a difference in soil moisture or herbivory’s association with fitness between habitat type.

Soil moisture availability is one of the main drivers of plant distribution and diversification (Stebbins 1952; Axelrod 1972). From our environmental analysis we learned that decreasing soil moisture was associated with decreasing survival on a fine spatial and temporal scale across all sites in our top linear mixed effects model (Table 1; Figure 2). However, a significant interaction between soil moisture, time, and habitat type demonstrated that soil moisture decayed and interacted with survival differently in granite outcrops vs. meadows (Table 1; Figure 2). In *M. guttatus*’ meadows soil moisture decayed in a slow linear fashion (Figure 2c&d), while in *M. laciniatus*’ granite outcrops soil moisture remained high and constant early in the season and then dropped precipitously mid-summer (Figure 2a&b). Different soil moisture regimes should select for different plant life histories. A fast cycling, drought escape strategy should be advantageous in the fast-drying granite outcrops while a longer lived, more water-use-efficient strategy should be advantageous in meadow habitat (Dudley 1996; Stanton et al. 2000; McKay et al. 2003; Anderson et al. 2011). The granite habitat’s water regime seems to have selected for *M*. *laciniatus’* rapid, early flowering life history.

We also found a difference in herbivory pressure between habitats with herbivory impacting survival on a fine spatial scale in *M. guttatus’* meadows, but not in granite outcrops (Table 1). There was also significantly less total herbivore damage in granite (Figure 3). Therefore it seems likely that herbivory acts as an important selective force in *M. guttatus*’ meadows, but not in *M. laciniatus’* granite outcrops. This is supported by the author’s observation of greater insect density in mesic meadows than in granite outcrops (K. Ferris personal observation). Perhaps *M. laciniatus* is not well adapted to herbivory which could contribute to the fecundity trade-off between *M. laciniatus* & *M. guttatus* in the meadow habitat. Herbivory is believed to be a major selective force driving plant diversification through coevolutionary forces (Ehrlich and Raven 1964; Janz 2011). In summary our data show that differences in soil moisture regime and herbivory between granite outcrops and nearby meadows likely contribute to divergent selection between *M. laciniatus* & *M. guttatus*’ habitats.

### Differential phenotypic plasticity between species

There is much debate in the literature about the role of phenotypic plasticity in adaptive evolution (Via et al. 1995; Schmitt et al. 1995; Dudley & Schmitt 1996; Ghalambor et al. 2007; Ghalambor et al. 2015). We discovered both phenotypic plasticity and genetic variation for plasticity (GxE) in flowering time, plant height, and flower size (Figure 5) in our reciprocal transplant. The presence of GxE in height, flower size, and flowering time indicates that phenotypic plasticity has the potential to respond to selection. Plasticity was in the expected direction, meaning in the same direction as species differences, for flowering time, plant height, and leaf shape. Plants in all genotypic classes (F_4_, *M. laciniatus, M. guttatus*) flowered earlier and were shorter in granite, and both *M*. *laciniatus* and F_4_’s were more highly lobed on average in granite than in meadow habitat (Table 2). *Mimulus laciniatus* had significantly greater flowering time plasticity than *M. guttatus* (Figure 5a). This ability to flower more readily in response to environmental cues may be advantageous given the rapid decreases in soil moisture and survival that characterize *M. laciniatus*’ native granite habitat (Figure 2a&b). On the other hand, the slow decline in soil moisture throughout the season in *M. guttatus*’ meadows would not require plants to flower as rapidly in response to drought (Figure 2 c&d). Therefore *M*. *laciniatus’s* granite environment might select for increased flowering time plasticity while *M. guttatus*’ meadows may not. Flowering time plasticity has been shown to be adaptive in other plant systems such as *Impatiens campensis* (Donohue et al. 2000; Donohue et al. 2001).

*Mimulus guttatus*, on the other hand, has greater environmentally induced plasticity in plant height than *M. laciniatus*. In a competitive environment like *M. guttatus’* meadows a plant’s ability to plastically change its height could be advantageous as a shade avoidance mechanism (Dudley & Schmitt 1996, Aphalo et al. 1999, Donohue et al. 2000, Weinig 2000). In a densely populated environment taller plants are able to outcompete other individuals for access to light and will therefore have higher photosynthetic rates and fitness (Schmitt et al. 1987, Aphalo et al. 1999). The significantly smaller size of *M. laciniatus* plants and leaves (Figure 5c&d) in meadows compared to the native *M. guttatus* suggests that *M. laciniatus* struggled to compete there. In the granite outcrop habitat *M. laciniatus* was only slightly smaller in stature and leaf size than *M. guttatus*. There was also selection for increased plant height in the meadow habitat, but decreased height in granite (Table 5). *Mimulus guttatus’* meadows have a much higher plant density than *M. laciniatus’* relatively depauperate granite outcrops (K. Ferris, personal observation), and small plants would be shaded out by other more vigorous species. To empirically test whether there is selection for increased flowering time plasticity in *M. laciniatus’* granite outcrops or height plasticity in *M. guttatus*’ meadows further experiments with replicate hybrid genotypes planted in each habitat would be necessary. Here we merely hypothesize that these differences in plasticity are advantageous given the known characteristics of each species’ habitat.

Another piece of evidence suggesting that *M. laciniatus* was at a competitive disadvantage in meadows was the direction of flower size plasticity between habitats. While *M. guttatus* displayed phenotypic plasticity in the expected direction, that is it had larger flowers in its native meadows than in the granite outcrops, *M. laciniatus* had larger flowers in granite outcrops than in meadows (Figure 5b). In our truncated Poisson distributed model larger flowers were selected for in both habitats, and consequently the decrease in *M. laciniatus’* flower size in meadows relative to granite suggests that *M. laciniatus* was at a competitive disadvantage in *M. guttatus’* habitat. Therefore competition with other species for light and resources could explain *M. laciniatus’* poor fecundity in *M. guttatus’* habitat and the species trade-off in lifetime fitness.

### Flowering time, plant size, and leaf shape contribute to differential habitat adaptation

We found that *M. laciniatus* and *M. guttatus* are differentially adapted to their respective habitats (Figure 1). *M. laciniatus* parents were significantly more likely to survive to flowering and produce more fruits in granite outcrops than *M. guttatus* parents (Table 1, Figure 4a). Interestingly in meadows we saw no difference in survival to flowering between species, but *M. guttatus* did produce more fruits than *M. laciniatus* in its meadow habitat (Figure 4b). A previous study also found that *M. laciniatus* had higher survival to flowering in granite than *M. guttatus*, but that the two survived equally well in meadows (Peterson et al. 2013). Many reciprocal transplant studies have found that one species or population does best in both habitats (Hereford et al. 2009) and this observation elicits the question, why doesn’t the population with highest overall fitness occur in both habitats? One potential reason for not observing a fitness trade-off in a reciprocal transplant study is that lifetime fitness was not measured. In this study we found an asymmetric result without a clear trade-off between species when we examined survival, but then found evidence of a species trade-off when we measured fecundity. The fact that we find selection against the non-native parent in each habitat indicates that there is ecological reproductive isolation between these species (Coyne & Orr 2004). *Mimulus laciniatus* occurs within the geographic range of *M. guttatus* (Ferris et al. 2014) and the two often co-occur within meters of each other creating the opportunity for gene flow (Ferris et al. 2016). Due to the lack of obvious intrinsic barriers between these species (K. Ferris personal observation) illustrated by the ease of crossing, this ecological component of reproductive isolation may be important for maintaining species boundaries in this system.

Our selection analysis found that earlier flowering time, small plant size, and lobed leaves underlie *M. laciniatus’* adaptation to its harsh granite outcrop habitat (Tables 4, 5, 6). Earlier flowering time is critical for drought escape in many annual plants that occupy seasonally dry environments (Kiang & Hamrick 1978; Fox 1990; Stanton et al. 2000; Mckay et al. 2003; Eckhart et al. 2004; Hall & Willis 2006; Franks et al. 2007; Anderson et al. 2012). This is because early flowering allows plants to reproduce before the rapid onset of seasonal drought in harsh, ephemeral habitats. Work with the herbaceous annual *Impatiens campensis* has found that early season drought selects for a rapid flowering, drought escape strategy, while late season drought favors slower developing genotypes with high WUE (Heschel & Reginos 2005). In continually moist environments we expect selection for later flowering or stabilizing selection on midseason flowering since later flowering plants are larger and more fecund (Mitchell-Olds 1996; Hall & Willis 2006; Anderson et al. 2012). *Mimulus laciniatus’* granite outcrop habitat dries out a month earlier in the growing season than nearby meadows where *M. guttatus* occurs (Figure 2; Peterson et al. 2013). In our reciprocal transplant earlier flowering significantly increased both the probability of a plant setting fruit and the number of fruits a plant was able to produce, in the granite outcrop habitat. Although we found evidence that selection acts more strongly on flowering time in granite outcrops than meadows (Tables 2&3), we did not detect selection for later flowering in the *M. guttatus* habitat. This may be because meadows dried out earlier than normal due to the 2013 California drought (Swain et al., 2014). Selection for earlier flowering has previously been detected in inland annual populations of *M. guttatus* (Hall & Willis 2006) indicating that this may be common in populations of this species with an annual life history.

We also detected differential selection on plant height between granite and meadow habitats. Plant size is intimately tied to plant fitness (Stearns 1992; Thornsberry et al. 2001; Falster & Westoby 2003). Larger plants are more fecund, but usually reproduce later in the growing season (Mitchell-Olds 1992; Mitchell-Olds 1996). In a seasonally dry environment later flowering places larger plants at risk of dying before reproduction. However, in a continually moist and densely populated environment larger plants have a competitive advantage both in terms of access to resources such as light and water and in increased fecundity due to their increased biomass (Mitchell-Olds 1992; Dudley & Schmitt 1996). In our reciprocal transplant we saw strong selection for taller plants in *M. guttatus’s* mesic meadow habitat both in terms of whether a plant produced fruit and the number of fruits it produced. In the seasonally dry granite habitat we also found that taller plants produced more fruit, but this selection was not as strong as in the meadow habitat. Importantly, we also found that smaller plants were significantly more likely to produce fruit in *M. laciniatus*’ granite habitat. Therefore we saw differential selection on plant height at first flower between species’ habitats in the direction of species’ differences. This indicates that plant height is an important component of differential habitat adaptation between *M. laciniatus* and *M. guttatus*.

Lobed leaves have been proposed to be adaptive in dry, marginal habitats because of their ability to be efficiently cooled by convection (Vogel 1968; Givnish 1978; Schuepp 1993; Nobel 2005) and because they have less drought stress prone tissue than entire leaves (Nicotra et al. 2011). However, it has been difficult to definitively demonstrate the adaptive significance of lobed leaf shape (Nicotra et al. 2011). Nicotra et al. (2008) demonstrated that lobed species of the South African genus *Pelargonium* have higher photosynthetic rates under high temperatures than their close entire-leaved relatives which they hypothesize is a strategy to maximize growth when water is available in seasonally dry habitats. However many traits are correlated when species diverge so it is difficult to say whether photosynthetic rates were a direct result of leaf shape or of evolutionary history. In the ivy leaf morning glory, *Ipomea hederaceae*, leaf shape has been shown to be under selection along a North-South cline (Campitelli & Stinchcombe 2013b) and in a segregating experimental field population (Bright & Rausher 2008). Northern populations of *I. hederaceae* are lobed, while Southern populations are predominantly entire-leaved (Campitelli & Stinchcombe 2013b). Our experiment is the first to directly test whether lobed leaf shape is under selection in a segregating inter-specific hybrid population. In the *M. guttatus* species complex lobed leaf shape is associated with the occupation of rocky, dry habitat (Ferris et al. 2015). Within the entire genus *Mimulus*, which consists of over 150 species, only two species, *M. laciniatus* and *M. filicifolius*, are fixed for lobed leaves and both specialize on granite outcrops (Ferris et al. 2014). Therefore we hypothesized that lobed leaf shape is an adaptation to this specific habitat in *Mimulus*. We are able to confirm this prediction in our reciprocal transplant where we found that lobed leaf shape was under positive selection in our F_4_ hybrid’s in granite, but not in meadow habitat (Tables 5&6).

We detected positive directional selection on flower size in both habitats (Table 6) in our Poisson regression, indicating this trait does not contribute to differential habitat adaptation between *M. guttatus* and *M. laciniatus* at this life history stage˙. In a study of *M. guttatus* inland annuals vs. coastal perennials, Hall and Willis (2006) also detected a pattern of selection for larger flowers across habitats. Increased flower size likely facilitates increased outcrossing via pollinator visitation, which would alleviate effects of inbreeding depression in both habitats. However, we did see evidence of differential selection on flower size between habitats in our binomial regression on whether a trait increased the probability of a plant producing fruit. The result was in the opposite direction of species differences since we observed selection for smaller flowers in the meadow habitat even though *M. guttatus* has larger flowers, while in granite larger flowers were more likely to fruit. Instead of being adaptive, the difference in floral size between the parental species may be due to selection on a correlated trait such as whole plant size or flowering time.

## Conclusions

Understanding the ecological variables and traits driving differential habitat adaptation is a key component of ecology and evolutionary biology, and is particularly relevant in light of the future increases in temperature and precipitation stress predicted to accompany global climate change. Adaptation is also critical for the generation and maintenance of biodiversity since differential adaptation between closely related sympatric species is an important component of reproductive isolation. We found that *M. laciniatus* and its supposed progenitor *M. guttatus* are differentially adapted to microhabitats that differ dramatically in drought regime and herbivory. Our results indicate that while early flowering time, rapid development, and lobed leaves are critical for plant fitness in *M. laciniatus’* harsh seasonally dry environment, traits such as herbivore resistance and large plant size may be more important in a competitive mesic environment like *M. guttatus’*. Our study is one of the few to find selection on lobed leaves, and the first to demonstrate that lobed leaves are adaptive in a seasonally dry environment. Additionally we find genetic variation for phenotypic plasticity between species in all traits indicating that plasticity can respond to selection in this system and may be a part of adaptation to each species habitat. By combining fine scale environmental surveys, large-scale reciprocal transplants, and phenotypic selection analysis we have reached a greater understanding of adaptation to a harsh marginal environment.

## ACKNOWLEDGEMENTS

We would like to thank Alex Gunderson for giving a month of his life to help the author drive cross-country and plant 3800 seedlings. The experiment could not have succeeded without his heroic efforts. We also thank Jenn Coughlan, Kathy Toll, and Annie Jeong for assistance collecting environmental data and censusing plants. Yosemite National Park provided permits and the Sierra Nevada Research Institute provided housing at the Yosemite Field Station. Funding was provided by an NSF DDIG (DEB-1210755), a California Native Plant Society Educational Grant, and a Sigma Xi Grants-in-Aid of Research awarded to Kathleen Ferris and John Willis, and by NSF En-Gen (EF-0723814) and LiT (IOS-1024966) grants awarded to John Willis.

## AUTHOR CONTRIBUTIONS

KF designed & performed the controlled crossing and field experiments, collected and analyzed all data, and wrote the first draft of the manuscript. JW helped with experimental design and provided feedback on the manuscript.

---

**Table S1.** Results of a logistic regression of survival to flowering on species and habitat to test for local adaptation. The model was Survival to Flowering ˜ Habitat (Granite or Meadow) + Species (*M. guttatus* or *M. laciniatus*) + Habitat*Species.

| Independent Variables | $\beta'$ | z-value | p-value |
| --- | --- | --- | --- |
| Plant(L) | 0.092 | 0.356 | 0.722 |
| Habitat(L) | 0.265 | 1.071 | 0.284 |
| Plant(L)*Habitat(L) | 0.964 | 2.876 | 0.004** |

**Table S2.**
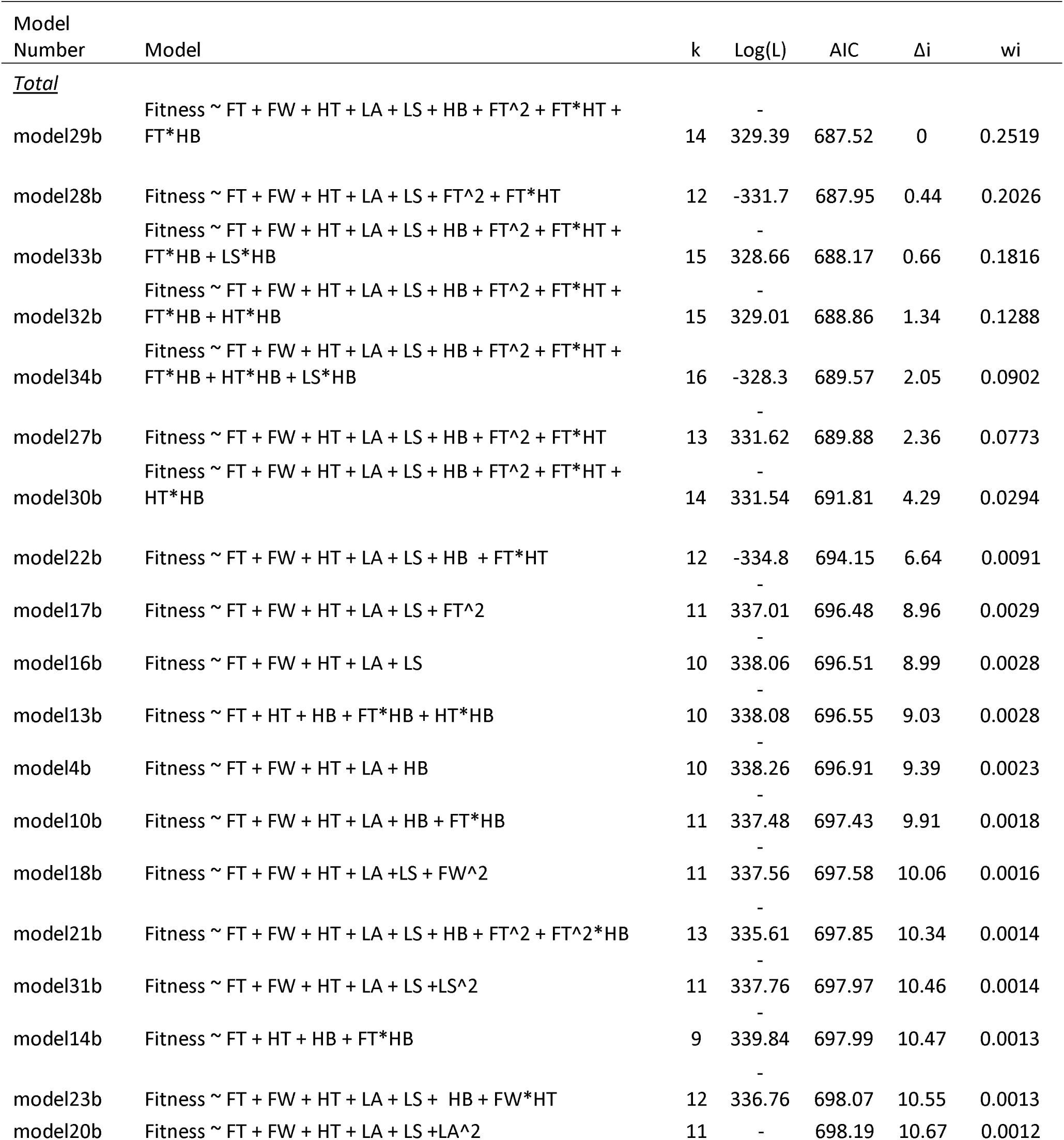
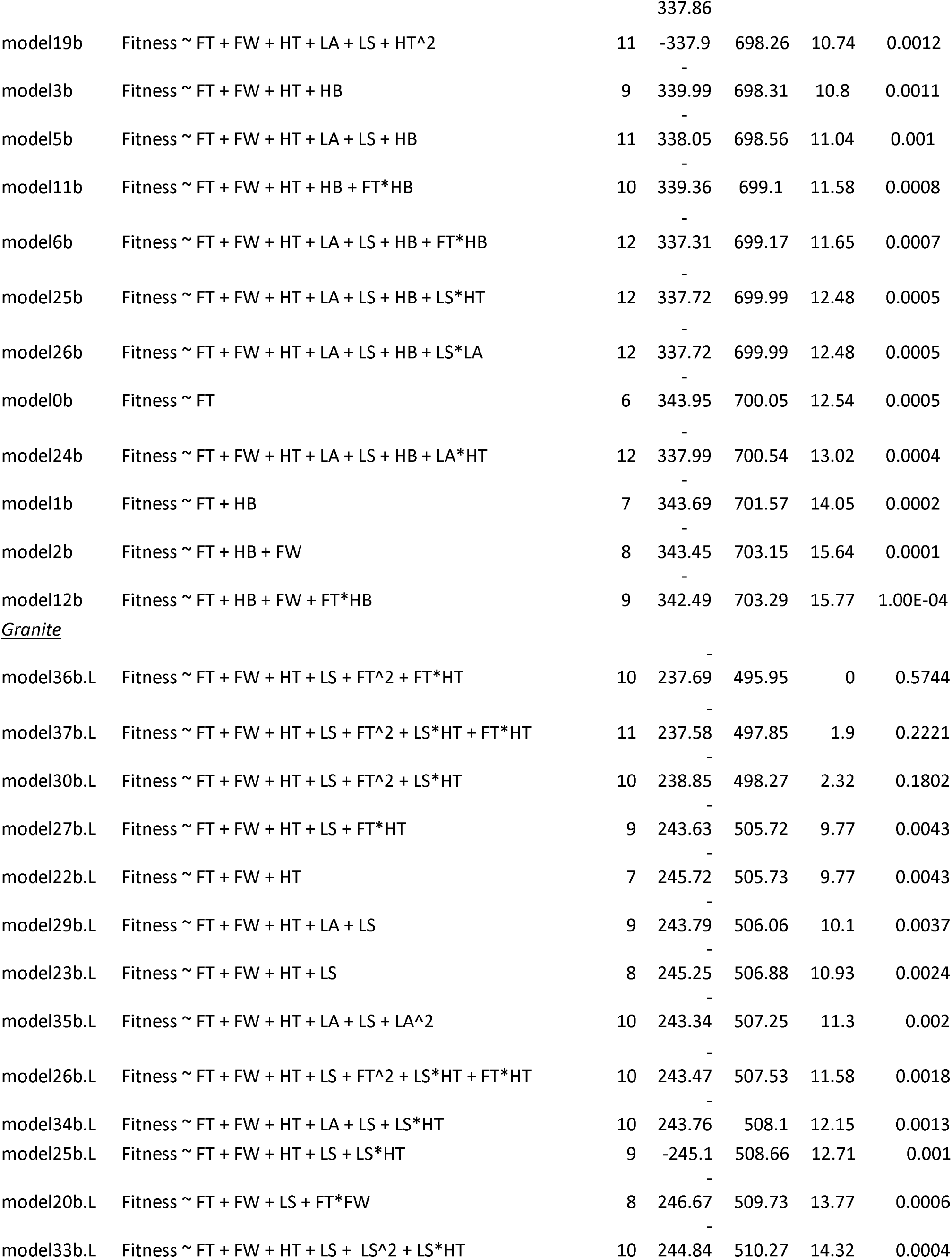
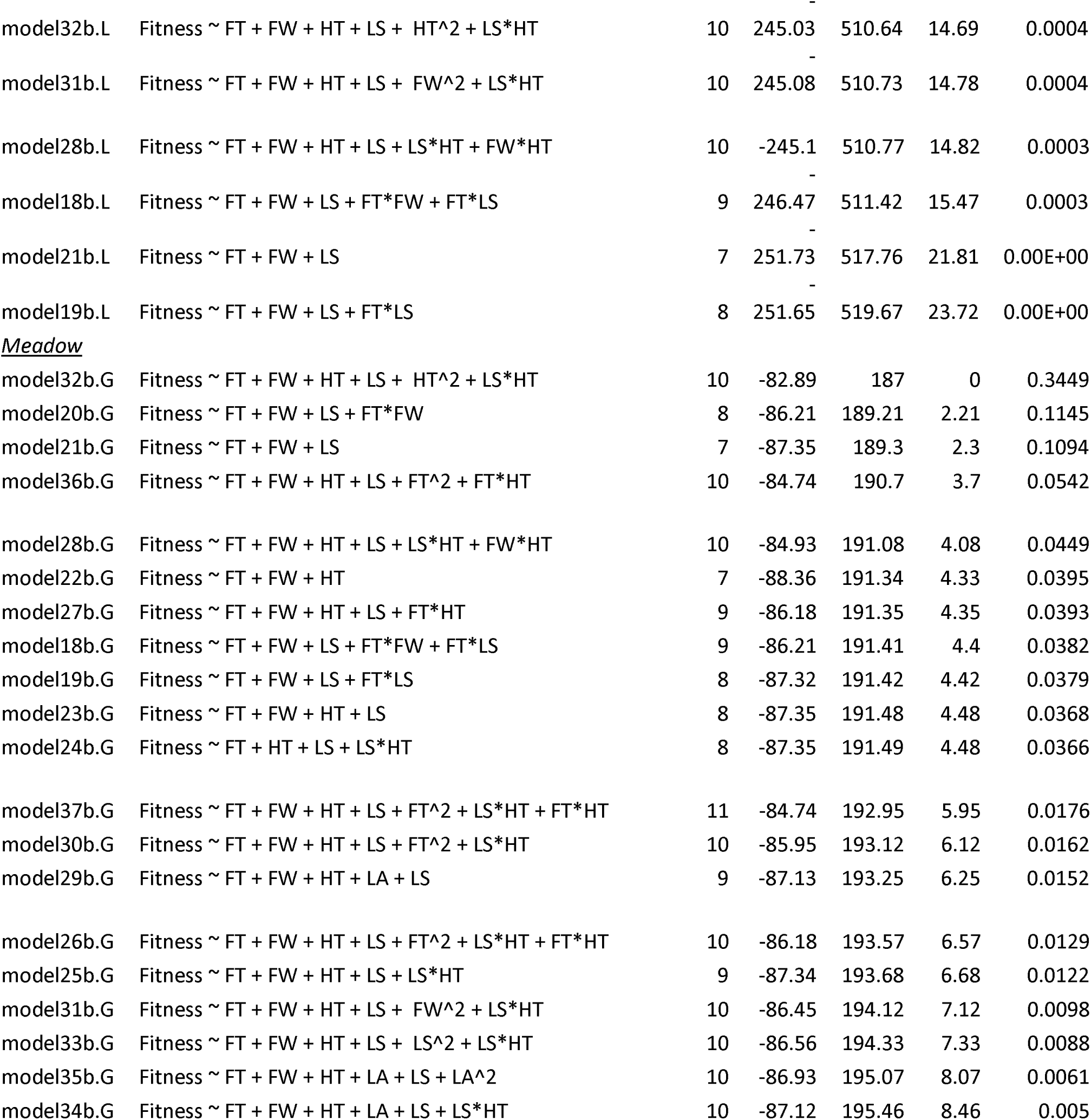
Complete list of all binomial distributed generalized linear mixed models testing which phenotypes influenced the probability that a plant produced fruit. For each we report the complete model, k which is the number of parameters, Log(L) which is the likelihood score, AIC score, Δi which is the difference between that model’s AIC and the top model’s AIC, and wi which is the model weight. FT stands for flowering time, FW stands for flower width, HT stands for height, LA stands for leaf area, LS stands for leaf shape, and HB stands for habitat.

**Table S3.**
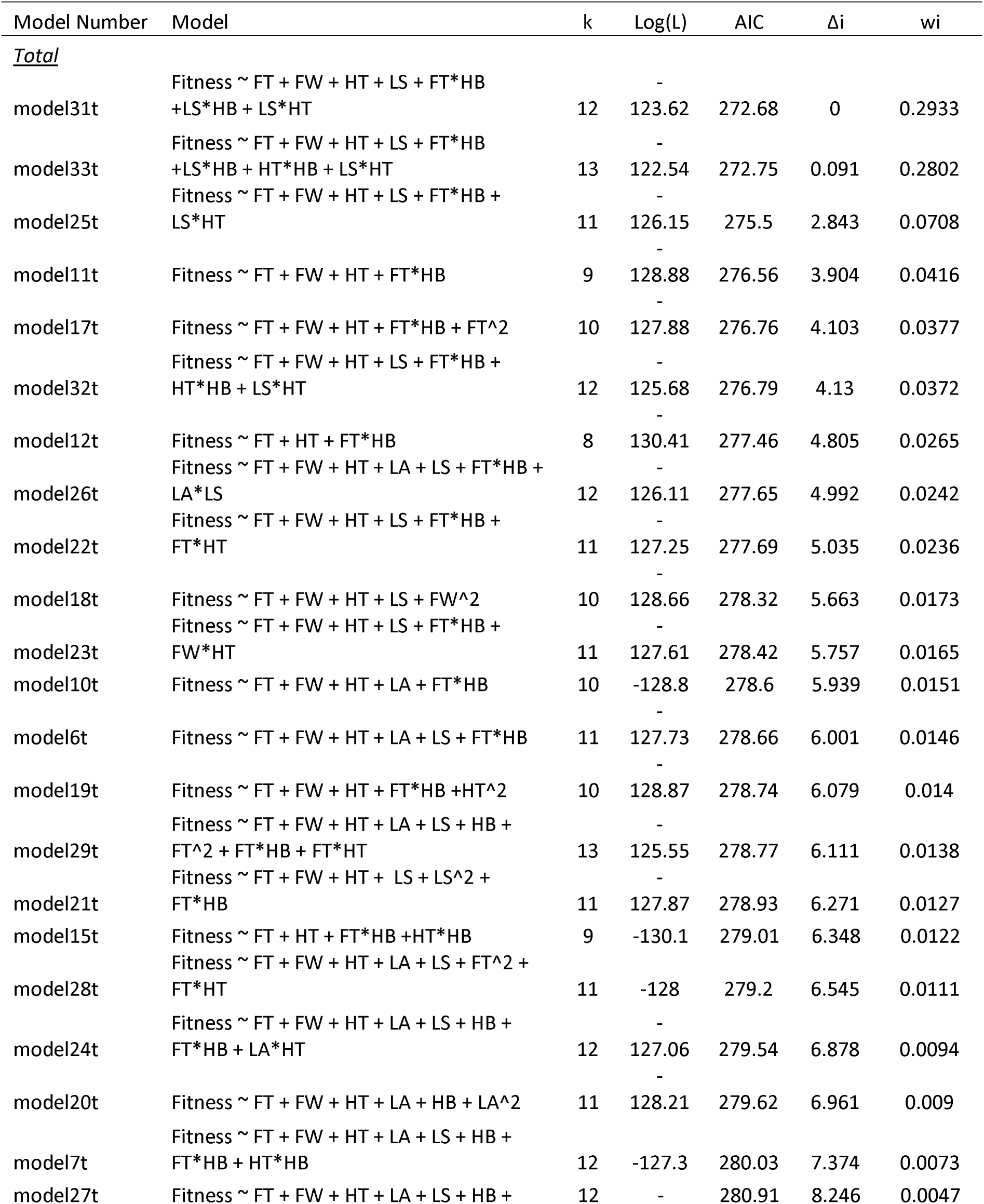
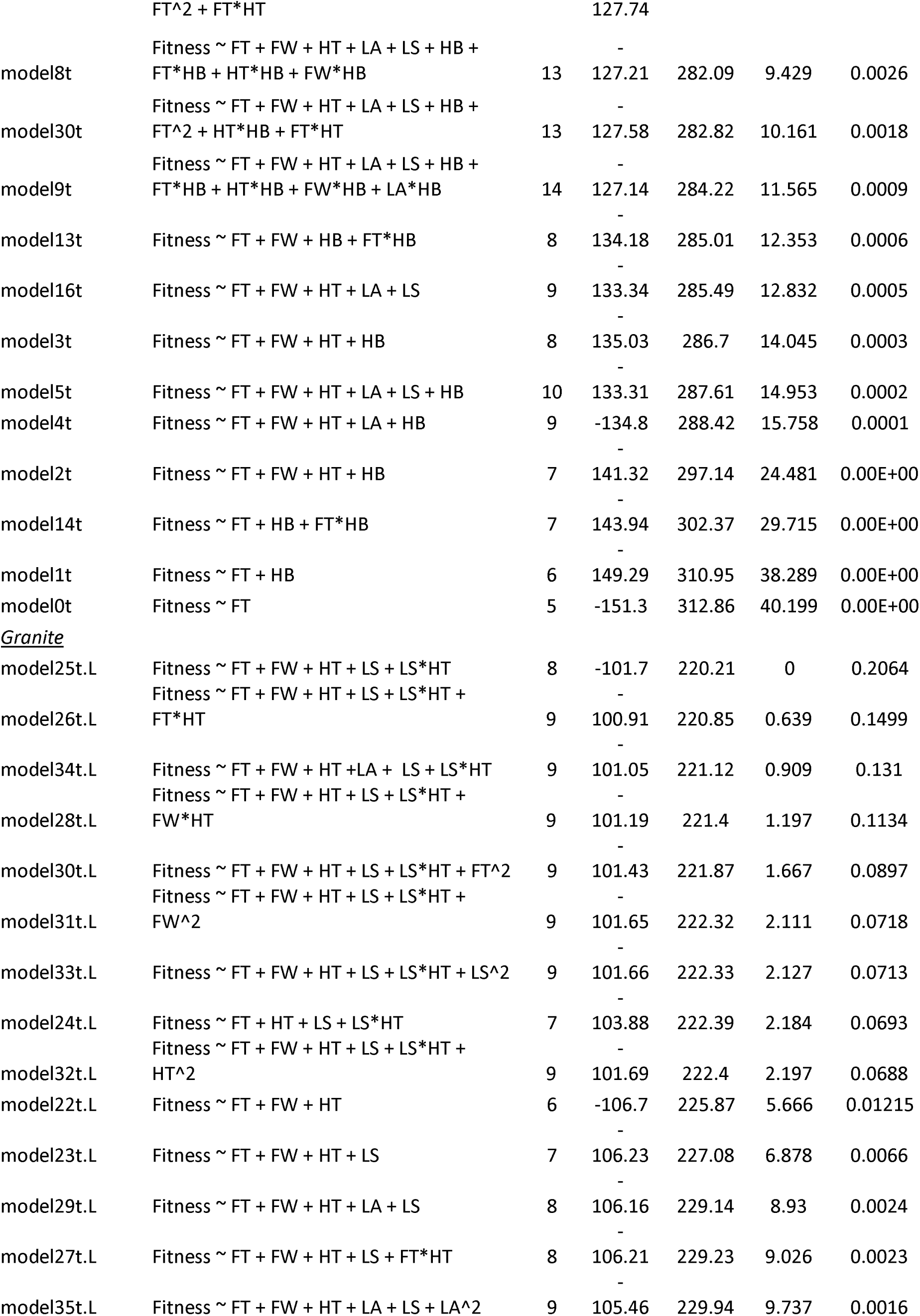
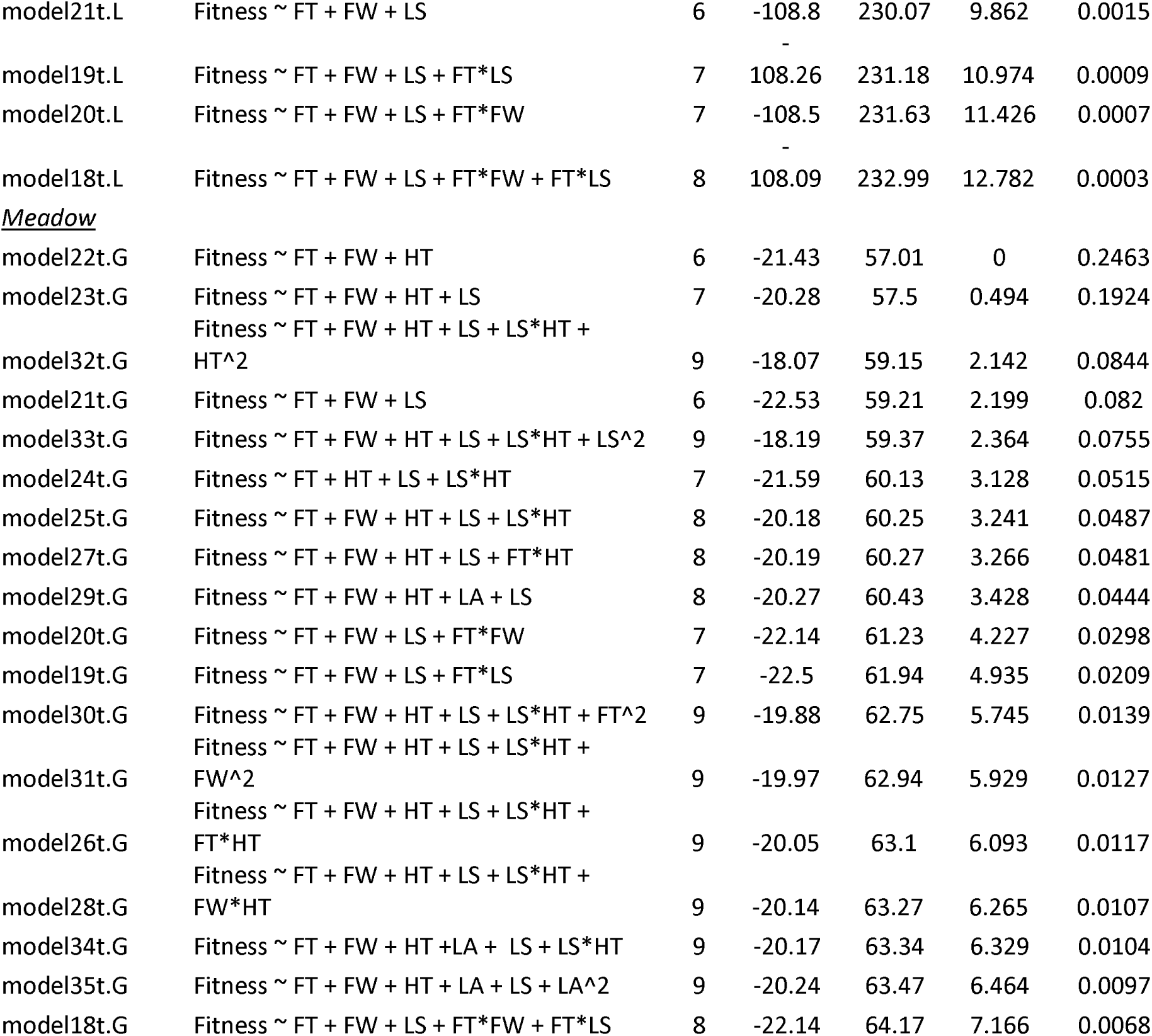
Complete list of all truncated Poisson distributed generalized linear mixed models testing which phenotypes influenced the number of fruits a plant produced. For each we report the complete model, k which is the number of parameters, Log(L) which is the likelihood score, AIC score, Δi which is the difference between that model’s AIC and the top model’s AIC, and wi which is the model weight. FT stands for flowering time, FW stands for flower width, HT stands for height, LA stands for leaf area, LS stands for leaf shape, and HB stands for habitat.

